# Comprehensive annotations of alternative splicing, lncRNAs, and circRNAs in seven species of flowering plants

**DOI:** 10.1101/2024.09.23.614400

**Authors:** Alexander Gabel, Christoph Schuster, Hajk-Georg Drost, Elliot M. Meyerowitz, Ivo Grosse

**Affiliations:** Institute of Computer Science, Martin Luther University Halle-Wittenberg, 06120 Halle (Saale), Germany; Institute of Medical Virology, Goethe-University Frankfurt, 60596 Frankfurt am Main, Germany; Infection Control and Antimicrobial Stewardship Unit, University Hospital Würzburg, 97080 Würzburg, Germany; Sainsbury Laboratory Cambridge, University of Cambridge, Cambridge CB2 1LR, United Kingdom; Computational Biology Group, Department of Molecular Biology, Max Planck Institute for Developmental Biology, Tübingen, Germany; Digital Biology Group, Division of Computational Biology, School of Life Sciences, University of Dundee, DD1 5EH Dundee, UK; Division of Biology and Biological Engineering, California Institute of Technology, Pasadena, CA 91125; Howard Hughes Medical Institute, California Institute of Technology, Pasadena, CA 91125; German Centre for Integrative Biodiversity, Research (iDiv) Halle-Jena-Leipzig, 04103 Leipzig, Germany

## Abstract

The successful occupation of terrestrial habitats by early plants was catalyzed by the adaptive evolution of new organs able to supply water and compensate for the gravitational impact under changing land conditions. Because plants are sessile, it has been proposed that the capacity of plants to innovate specialized organs is driven by a complex interplay between developmental gene expression and regulatory RNAs such as lncRNAs and circRNAs, evolving through sequence polymorphisms, genomic rearrangements, and expression divergence. Despite the importance of alternative and noncoding transcripts for enabling plant life, their accurate quantification causes major challenges as many of them are expressed at low levels and tissue-specific. Here, we describe the re-annotation of seven land plant genomes based on deep, organ-specific, ribo-depleted developmental RNA-Seq data. Using this comparative resource, we uncover 5,000 new lncRNAs and 2,000 new circular RNAs that are promising candidates for further functional investigation. Our annotation will become a reference catalog for studies on plant organ evolution and for uncovering tissue-specific patterns of transcript emergence.

## Introduction

Unveiling the functional mechanisms driving land plant adaptation is key to predicting the survival capacity of plants during our time of global climate change [1]. Next to protein-coding genes, the role of long non-coding (lncRNAs) and circular RNAs (circRNAs) in driving tissue-specific gene expression has emerged as a promising direction for functional studies in recent years. LncRNAs are a class of RNAs with a sequence length of at least 200 nucleotides and no apparent protein-coding function. However, some lncRNAs encode for small proteins [2] and were proposed to serve as initial source during the origination of new protein-coding genes [3].

The regulatory function of lncRNAs has been demonstrated in different species at the tissue and organ level with primary focus on animal examples [4–7], but plant lncRNAs also seem to be involved in similar functions [8–11]. Plant focused transcriptome-wide studies show that lncRNAs are involved in regulating the transcription machinery, histone modifications, alternative splicing, miRNA activity, and protein-protein interactions [12].

In addition, circRNAs are a class of RNA molecules that originate from a circularization event, called backsplicing, which is defined by a downstream 3’ splicing donor site and a 5’ upstream splicing acceptor site (Supplementary Fig. 3a). Complex tissue- and developmental stage-specific expression of circRNAs suggests that their role is associated with post-transcriptional gene regulation [13–17].

The functional study of lncRNAs and circRNAs in land plants remains underexplored and mostly restricted to only a few model organisms such as *Arabidopsis thaliana* (TAIR [18] Araport [19], PLncDB [20], and at population scale [21]). For a wide range of other species, however, the genome-wide identification of developmental genes, lncRNAs and circRNAs remains incomplete and often out of reach.

Studies in animals [5, 7] and plants [19] have shown that lncRNA and circRNA expression is often strongly organ-dependent, suggesting that basing annotation derived on a broad set of organs may be required to understand the true abundance of expressed genomic loci. We therefore sourced our comprehensive organ-specific, ribo-depleted total RNA-Seq gene expression atlas of seven flowering plants [22] and computed de novo annotations of splice variants, lncRNAs, and circRNAs for all seven plants. To enable this comprehensive annotation task, we developed a dedicated Snakemake [23] workflow that allowed us to standardize the quantification of species- and organ-specific protein-coding transcripts, lncRNAs, and circRNAs (Figure 1). Using our computational workflow, we characterized all newly discovered RNA molecules and prepared a comparative annotation for seven flowering plants.

**Figure 1.**
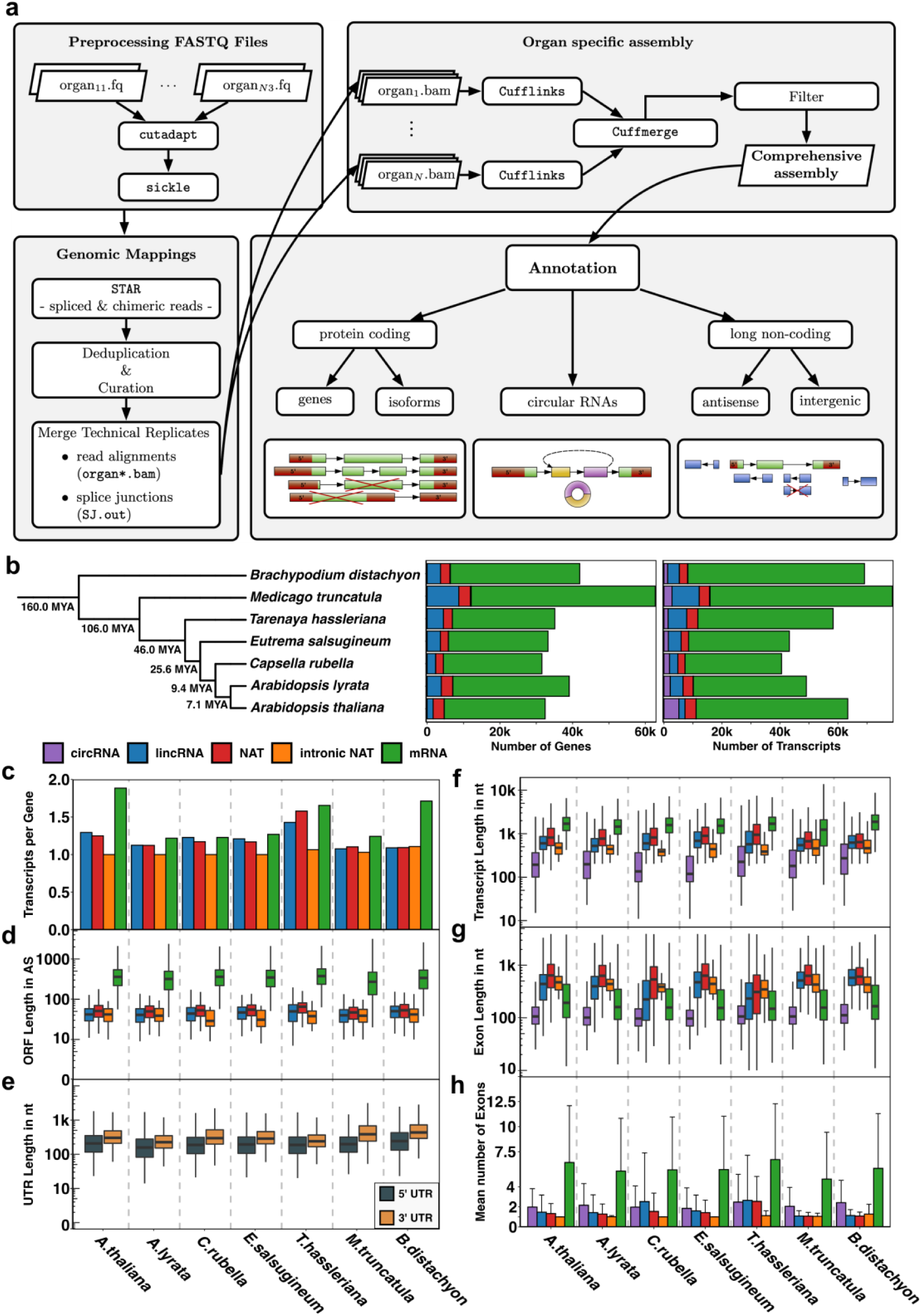
Overview of the DevSeq annotation workflow and quantification of protein coding genes, lncRNAs, and circRNAs for seven flowering plants. a, The first part of the pipeline summarizes the preprocessing step which includes adapter clipping and quality trimming of the raw RNA-Seq data in fastq-format. In the next processing step the post-processed RNA-Seq reads are aligned against the reference genome of the respective species using the mapper STAR [54], followed by read-deduplication and merging of technical replicates. Subsequently, Cufflinks [55, 56] was employed to assemble organ-specific transcriptomes for each flowering plant species separately. The organ-specific transcriptomes were merged within each species to achieve a comprehensive assembly. Low-quality transcripts were removed using stringent filter criteria. The final assembly then served as the basis for predicting isoform sensitive protein-coding loci, circular RNAs, and long non-coding RNAs. b, Quantification of the number of genes and transcripts that were found using the DevSeq annotation workflow. Species are ordered according to their taxonomic distance from the reference species A. thaliana. Estimated divergence times in million years ago (MYA) from TimeTree of Life [57]. c - h, Further output of the DevSeq annotation workflow quantifying for each flowering plant species independently and divided by circRNAs, lincRNAs, NATs, intronic NATs, and mRNAs c, Number of transcripts per gene, d, Open Reading Frame (ORF) length, e, UTR length, f, Transcript length, g, Exon length, h, Average number of exons (arithmetic mean).

## Results

Our genome annotation workflow (Figure 1a) is designed to enable a species-specific comparison of protein-coding transcripts, long non-coding RNAs, and circular RNAs of *A. thaliana*, *Arabidopsis lyrata*, *Capsella rubella*, *Eutrema salsugineum*, *Tarenaya hassleriana*, *Medicago truncatula*, and *Brachypodium distachyon*. For each plant species, we sourced sequencing data from root, hypocotyl (mesocotyl from *B. distachyon*, respectively), leaves, vegetative apex, inflorescence apex, flower, stamen, carpel and the mature pollen, consisting of two cell types. Since the workflow utilizes organ-specific developmental RNA-Seq data to create comprehensive species-specific annotations, we refer to this workflow as the DevSeq annotation workflow and the resulting annotation catalog as DevSeq annotation. For each species, the workflow reconstructed full-length transcript models of all three RNA species (mRNA, lncRNA, and circRNA). For lncRNAs, we defined further subgroups: long intergenic non-coding RNAs (lincRNAs), natural antisense long non-coding transcripts (NATs), and intronic natural antisense long non-coding transcripts (intronic NATs).

Because transcript isoforms and lncRNAs are often expressed in an organ-specific manner [5, 7, 24–27], we extended our analysis by reconstructing the transcripts for each organ. For each species, we merged the organ-specific transcriptome assemblies to a comprehensive assembly representing the expressed transcripts found in the sequenced organs of each plant. Based on the assembly and the genomic mapping, we classified the assembled transcripts into protein-coding, lncRNAs (and their subgroups), and circRNAs. Figure 1b shows the absolute frequencies of annotated genes and transcripts for all seven plant species.

We validated the annotation completeness by performing sequence similarity searches against the Benchmarking Universal Single-Copy Orthologs (BUSCO) [28] and against the curated set of embryophyta-specific single-copy orthologs, which ideally should be present in all analyzed plant species. The analyses showed that between 96.4% and 99.5% of the curated BUSCO sequences were found (Supplementary Table 1). For all species but *E. salsugineum* and *M. truncatula* the DevSeq annotation approached apparent completeness.

We compared our DevSeq annotation against the currently published reference annotation of each species. Based on this overlap assessment with previously reported (expressed) loci, we identified novel protein-coding isoforms and novel putative protein-coding loci ranging from 126 (*A. thaliana*) to 1,256 (*B. distachyon*) loci (Supplementary Table 2).

As a result, our DevSeq annotation contains thousands of novel and previously unreported protein-coding transcripts (Supplementary Table 3). Due to the high sequencing depth of our RNA-Seq libraries and the broad variety of organs, we were able to determine poorly expressed and previously neglected organ-specific transcripts. In addition, we verified previously published isoforms and elongated their 5’ and/or 3’ boundaries when our transcription/isoform evidence was in support of this reannotation (Supplementary Table 3). Furthermore, our DevSeq workflow discovered between 2,800 (*T. hassleriana*) and over 6,000 (*B. distachyon*) novel isoform transcripts.

While updating the previously published set of annotated transcripts, we detected a systematic reason for the increase in the number of splicing events in protein-coding isoforms. In detail, the fraction of alternative 3’ ends slightly increases in most species, except for *A. lyrata* and *E. salsugineum* (Fig. 2a). We identified the extension of alternative 3’ ends as the main alternative splicing event within the published reference annotations as well as in our new DevSeq annotation. Next, we found alternative 5’ ends and intron retention events to be the second largest fraction of alternative splicing events. In our DevSeq annotation the total number of alternative 5’ ends increased in all species compared to the published reference annotations. This effect could be explained by random priming during RNA-Seq library preparation. Previous work has shown that random priming leads to more uniformly distributed reads in genomic mapping but may also introduce coverage biases at 5’ ends [29]. We therefore recommend caution when interpreting the excess of alternative 5’ ends.

**Figure 2.**
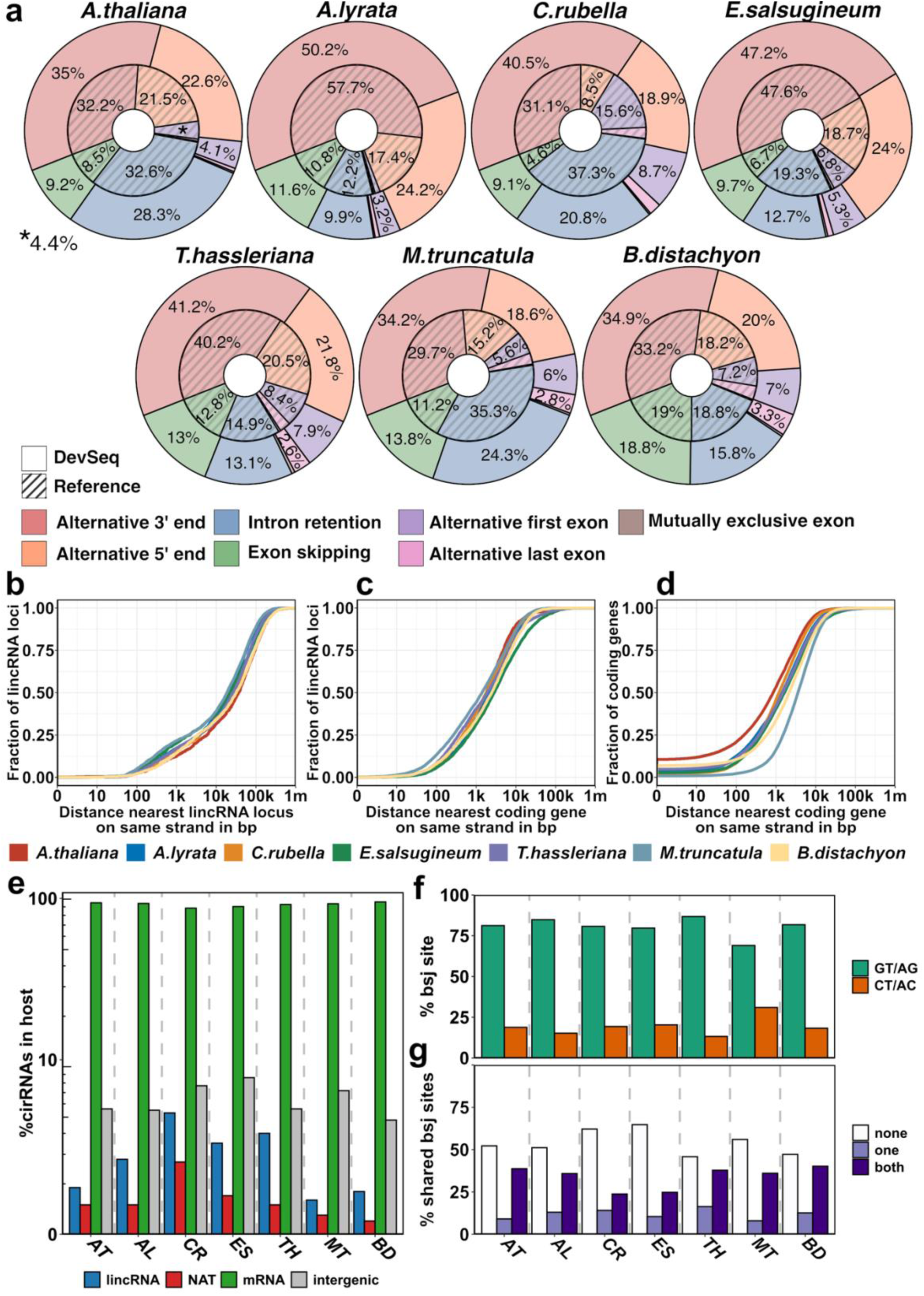
Summary statistics and genomic characterization of coding and non-coding transcripts. a, Direct comparison of our *de novo* annotated splicing events (DevSeq) against the published reference annotation currently employed in each respective species. The outer circles denote the fraction of splicing events generated using our DevSeq annotation workflow. The striped inner circles represent the proportions of splicing events observed in the respective reference annotation. b - d, Genomic distances of b, lincRNAs with respect to their neighboring lincRNAs, c, protein-coding genes with respect to their nearest coding gene on the same strand, and d, protein-coding genes to their neighboring protein-coding genes. e, Percentage of circRNAs hosted within other RNA molecules >200 nt or within intergenic regions. f, Fraction of back-spliced junction sites (bsj, donor/acceptor) of circRNAs. g, Fraction of back-spliced junction sites (bsj) shared with host protein-coding transcript.

Due to the experimental design of the total RNA sequencing data obtained for this study, we could not distinguish between introns retained after splicing of a pre-mRNA transcript and the primary transcript. To reduce potentially false annotations, we discarded all novel protein-coding isoforms that showed evidence of origin from retained introns. In contrast, isoforms from the reference annotations showing intron retention events are integrated into our DevSeq annotation. The alternative 3’ and 5’ ends represent the largest fractions of novel splice variants (Fig. 2a). To verify that our annotation has not introduced a technical bias when annotating transcriptomes, we compared the lengths of the observed 5’ and 3’ untranslated regions (UTR) across all seven species (Fig. 1e). This comparison of UTRs (Fig.1e) illustrates that in all species 5’ UTRs were shorter than 3’ UTRs, which is a well-known feature of eukaryotic mRNAs [26]. Comparing the UTR lengths between all species, we observe that the majority of plants follow a similar distribution of UTR length. Only *M. truncatula* and *B. distachyon* showed an increased overall length in 3’ UTR sequences.

Next, we classified assembled transcripts without any overlap with annotated protein-coding genes on the same strand as potential candidates for lncRNAs. After calculating their coding potential [30] and performing several filter steps (Methods), we identified thousands of lncRNAs (Fig. 1b) for all seven plant species.

Additionally, we compared the average number of isoforms that were found in lncRNAs and protein-coding transcripts (Fig. 1c). Except for *T. hassleriana*, lncRNA loci transcribe (on average) between 1.00 - 1.25 transcripts per isoform. This low number of transcripts per lncRNA locus was also reported by other studies [31–35]. In contrast to these studies, we observed that *A. lyrata*, *C. rubella*, *E. salsugineum*, and *M. truncatula* yield comparable numbers of transcript isoforms for protein-coding genes. Only *A. thaliana* and *B. distachyon* were exceptions, showing an increase of transcript isoforms in protein-coding genes comparable to lncRNA loci.

We found the median transcript length of lncRNAs to be approximately 600 nt, which is an order of magnitude shorter than the median length of protein-coding genes at approximately 2000 nt (Fig. 1f). Within lncRNAs, NATs showed the largest transcript lengths followed by lincRNAs, intronic NATs, and circRNAs.

The pattern of transcript length distributions of the RNA species was consistent in all plant species with small exceptions regarding the transcript length of protein-coding transcripts from *M. truncatula*. The median transcript length of protein-coding transcripts was similar compared to the other plant species, but the interquartile range (IQR) varied drastically, with longer sequences reaching over 10,000 nt.

Exon length distributions were also similar for the majority of species except for *T. hassleriana* showing a very broad distribution of exon lengths for lincRNAs and NATs (Fig. 1g). Regarding the distribution of exon lengths (Fig. 1g) and the mean number of exons per transcript (Fig. 1h) over the different RNA species, lncRNAs seemed to have long exons with a median of 500 nt, but contained on average only 1.5 exons per transcript. In *C. rubella* and *T. hassleriana* the number of exons per transcript was around three and the exon length distributions of lncRNAs were similar to protein-coding genes. The similarity might reflect limitations in the prediction of lncRNAs, which seems to be in contrast to the ORF lengths (Fig. 1b). This result provides further evidence that lncRNAs may serve as sources for novel peptides [3].

The classification of lncRNAs depended on their genomic location with respect to protein-coding loci. We found that in all species, a substantial fraction of lincRNA loci is within 500 nt from their neighbors, which could implicate a possible clustering of lincRNAs along some chromosomes (Fig. 2b). However, over 75% of lincRNA loci were fairly distant from other lincRNA loci, with observed distances of up to 5,000 nt. In comparison (Fig. 1c), the median distance of lincRNA loci to the closest protein-coding genes was just 1,000 nt. The maximal distance to protein-coding genes increased to over 50,000 nt. Such loci were previously described as isolated lincRNA loci [35].

On the other side of the distance spectrum, we determined lincRNA loci in close proximity between 0 and <1,000 nt. We classified lncRNA loci as intergenic if their start or end coordinates did not overlap with protein-coding genes [2, 36] or if they were not within a distance between 500 nt and 1,000 nt to a protein-coding gene [27, 37–39]. The later definition of lincRNAs is often combined with the notion of bidirectional lncRNAs, which are thought to share the same promoter region with a neighboring protein-coding gene. To our knowledge, there is no comprehensive definition of bidirectional lncRNAs that could be applied to plants and animals interchangeably. From the differences we observed between animal and plant bidirectional lncRNAs, we envision that a universal definition of bidirectional lncRNAs would need to depend on the location of the promoter to the closest protein-coding gene.

The DevSeq annotation workflow also detected thousands of novel circRNAs. Predominantly, we found that circRNA loci often overlap with protein-coding loci, i.e., 87-95% of all observed circRNAs were located within protein-coding genes (Fig. 2e). We detected ∼4-8% circRNAs within intergenic regions and 1-5% within lncRNAs (Fig. 2e).

The observed transcript length of circRNAs ranged between 10 nt and 3,000 nt (Fig. 1f). The median length was 200 nt and thus much shorter than the median transcript length of lncRNAs or protein-coding transcripts. In contrast, circRNAs seem to contain more exons than lncRNAs (Fig. 1h).

The majority of back-splicing event forming a circRNA were GT/AG donor-acceptor pairs. However, GT/AG donor-acceptor sites denoted the largest fraction of transcript splice sites across all RNA types and was consistent across all seven plant species (Fig. 2f). About 10-25% of all back-splice events contained CT/AC donor-acceptor sites. We also found that the back-splice events that were not intergenic generally shared no splice site with their host transcript (Fig. 2g) or shared both splice sites with its linear host transcript. A minority of back-splice events shared only a donor or an acceptor splice site with their linear host transcript.

The GC content is a characteristic genomic feature to differentiate between coding and non-coding regions. Grasses were shown to contain higher GC contents than other angiosperms [40–43] with most evidence derived from quantifying GC abundance and GC variation between intergenic and protein-coding regions [41, 43–46].

Comparing intronic and exonic sequences (Supplementary Fig. 4), we found that in all RNA species, the GC content was lower in intronic regions than in exonic regions. This low GC content is consistent with other observations from protein-coding regions in plants and animals [47–49].

*B. distachyon*, as a representative of monocot grasses showed the highest GC content and the widest range of GC abundance in all four RNA species (Fig. 3 a - d; Supplementary Table 5). We further categorized the transcripts of each RNA species within each plant into five groups based on their transcript length (Fig. 3; Supplementary Table 5). The IQR decreased in the majority of RNA species and the standard deviation of GC content increased with transcript lengths whereas the median of GC content is constant.

**Figure 3.**
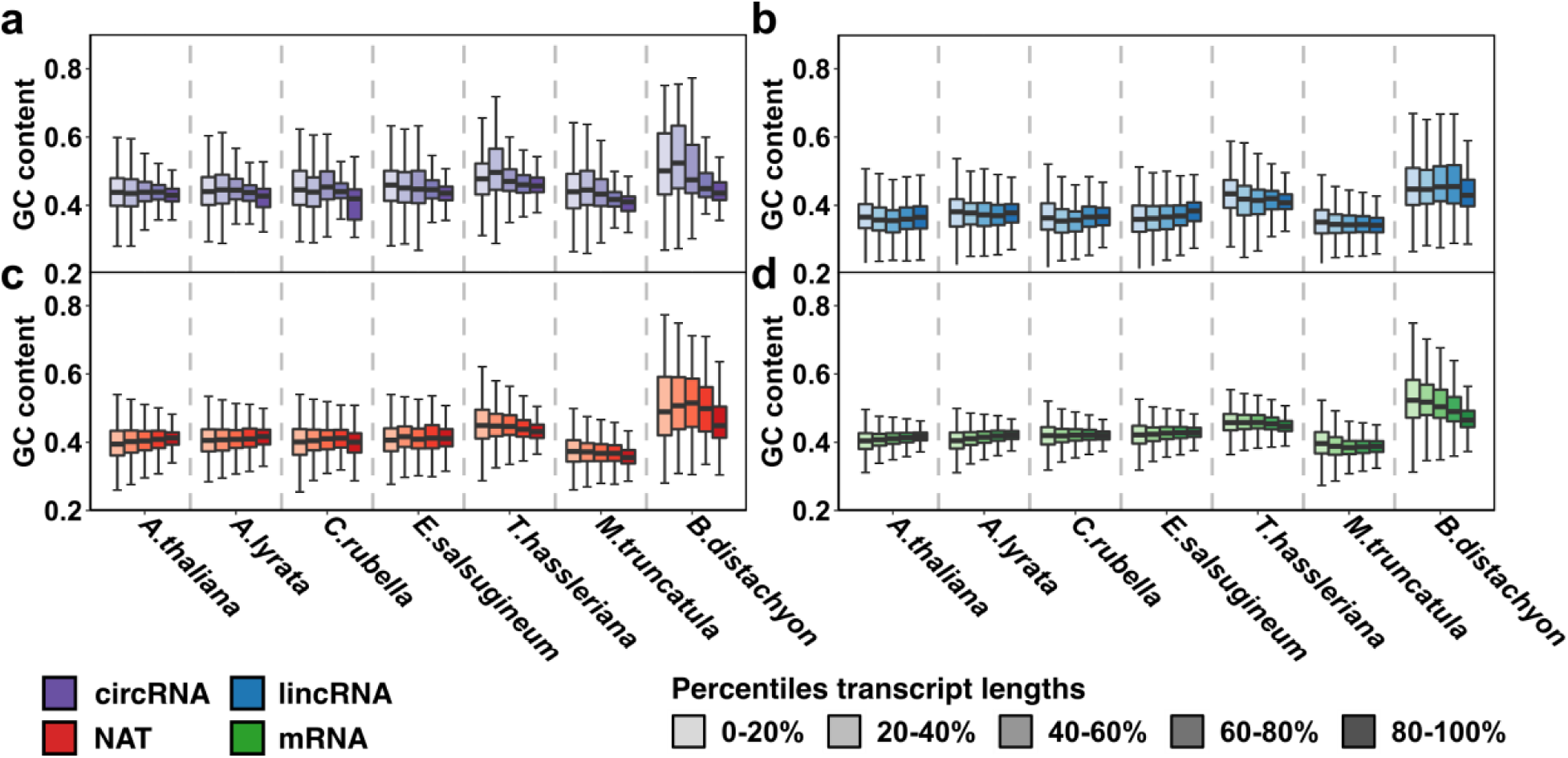
GC content vs. transcript lengths. (a) CircRNAs, (b) lincRNAs, (c) NATs, and (d) protein-coding mRNAs of each plant species are grouped into equally sized quintiles by their transcript lengths. For each group, we plot the relative GC content of each transcript within each quintile. Supplementary Table 5 presents detailed information on transcript lengths and GC content within each quintile.

While *B. distachyon* showed the highest GC content in all four RNA species, it is the only plant species in which the GC content decreased while the mRNA transcript length increased (Fig. 3d). A detailed investigation of GC contents of coding and non-coding transcripts in flowering plants was not the focus of this study and will be part of future work.

## Conclusions

In this study, we introduced the DevSeq annotation workflow to predict and annotate protein-coding genes, splice variants, lncRNAs, and circRNAs in seven flowering plants based on total RNA-Seq data conducted in an organ-specific manner. The fully reproducible Snakemake [23] workflow relied on genomic reference sequences and annotations.

For the majority of our plant species, no prior annotation of lncRNAs existed. We tailored the prediction of lncRNAs for each plant species based on their annotated protein sequences. Overall, we detected approximately 5,000 novel lncRNAs and 2,000 novel circular RNAs for each plant species, identified thousands of novel protein-coding transcript isoforms, and updated currently annotated transcripts by elongating their 3’ and 5’ ends.

The seven annotations serve as a resource for organ-specific annotations of noncoding RNAs in plants and could potentially deepen our understanding of the developmental transcriptomes in complex plant species. Notably, we used the annotations described in this work as a starting point for transcript quantification to investigate the evolution of organ-specific transcriptomes in flowering plants [22]. We sought to shed light on the non-coding landscape of divergent land plant species, with a particular focus on their contribution to organ formation and tissue specific function.

We envision that our DevSeq annotations catalog will serve as a community resource for functional screening with a particular focus on organ-specific expression of non-coding RNAs. Our standardized workflow allows direct comparison of annotation results between plant species. We learned that a broad RNA library sampling from diverse plant organs with a fine developmental resolution can unveil a significant proportion of new lncRNAs, circRNAs, and splice variants with putative regulatory roles in organ formation. Finally, our reannotation of seven plant genomes provides a comparative framework for studies interested in the evolutionary processes shaping non-coding regions and their functional co-option for regulating organ formation and phenotypic innovation.

## Author contributions

A.G., H.G.D., C.S., E.M.M., I.G. conceived the study; A.G. implemented the DevSeq annotation workflow and generated annotations with input from C.S., H.G.D, E.M.M., and I.G.; A.G., C.S., H.G.D. analyzed and interpreted data. A.G., H.G.D., and C.S. drafted the manuscript with contributions from E.M.M. and I.G.

## Acknowledgements

We thank Jan Grau, Ioana Lemnian, Martin Porsch, and Claus Weinholdt for valuable scientific discussions, and for financial support we thank the German Research Foundation (DFG INST 271/339-1 FUGG, FZT 118, GR 3526/2, GR 3526/6, GR 3526/7, and GR 3526/8 to I.G.) and the Gatsby Charitable Foundation (GAT3731/DAA to E.M.M.). The E.M.M. lab is also supported by the Howard Hughes Medical Institute. H.G.D. in particular thanks Detlef Weigel for ongoing support and sponsorship of his work and for the support by the Max Planck Society. A.G. and H.G.D. thank the Stickstoffwerke Piesteritz for the SKWP Research Prize 2013 and 2017.

## Methods

### Sample collection and RNA-Seq library preparation

Total RNA-Seq data was generated from seven plant species with an evolutionary divergence ranging from 7.1 million years ago (MYA) up to 160 MYA as representatives of the flowering plants phylogeny. The ecotypes are *Arabidopsis thaliana* (Col-0), *Arabidopsis lyrata* (MN47)[58], *Capsella rubella* (Monte Gargano) [59], *Eutrema salsugineum* (Shandong) [60], *Tarenaya hassleriana* (ES1100)[61], *Medicago truncatula* (A17) [62], and *Brachypodium distachyon* (Bd21) [63]. All chosen species possessed complete or partially complete genome sequences, and the annotations of each species contained at least information about protein-coding genes.

We used various tissues and developmental stages for *A.thaliana*, and root, hypocothyl (mesocotyl for *B. distachyon*, respectively), leaf, apex vegetative, apex inflorescence, flower, stamen, carpel, and pollen for the remaining plant species. In total, we generated 303 samples [22].

A detailed description on sample collection and RNA-Seq library preparation is described in [22]. Briefly, the sequencing libraries were prepared using the Illumina TruSeq Stranded Total RNA with Ribo-Zero Plant preparation kit. The resulting sequencing libraries were sequenced with 75bp paired-end reads on an Illumina HiSeq4000, except one lane of re-sequencing on Illumina NextSeq500.

Sequencing of total RNA allowed capturing transcripts with poly(A) tails (poly(A^+^)) and without poly(A) tails (poly(A^−^)). This was essential for annotating novel lncRNAs transcribed by RNA polymerase IV and V [64–67] and for circular RNAs [15, 16, 68] lacking polyadenylated 3’ ends. All samples were sequenced with additional ERCC RNA Spike-In control Mix 1 [69] allowing the definition of an objective expression cutoff to differentiate lowly expressed transcripts from technical noise by relating the concentration of ERCC Spike-Ins to their subsequently estimated expression values. Hence, the expression values of ERCC Spike-ins with low concentrations served as a lower bound to call a particular transcript as expressed.

### Read mapping and transcript assembly

Read quality was assessed by *fastqc* (version 0.11.5) [70] to evaluate possible sequencing errors, contaminations during library preparation and/or subsequent sequencing. Sequencing adapters were removed by *cutadapt* (version 1.10) [71]. Quality trimming of reads was performed by sickle (version 1.33) [72] with parameters pe −t sanger −q 20 −l 50.

Paired-end reads were mapped against their respective reference genome using the RNA-Seq aligner STAR (version 2.7.1a) [54] with the parameters --alignIntronMin 20 --alignIntronMax 10000 -- bamRemoveDuplicatesType UniqueIdenticalNotMulti --chimSegmentMin 10 --chimOutType Junctions WithinBAM --outFilterMismatchNmax 3 --outFilterMultimapNmax 1.

Genomic reference sequences and annotations for the genome-guided transcriptome assembly and subsequent annotation procedures were obtained from Ensembl [73], release 34 (for *A. thaliana*), Phytozome [74], version 12 (for *A. lyrata*, *C. rubella*, *E. salsugineum*, *M. truncatula*, and *B. distachyon*), and RefSeq [75] (for *T. hassleriana*). Detailed information is listed in Supplementary Table 4.

After read mapping, a duplication analysis was performed to detect overrepresented fragments, which are characteristic for PCR artifacts. Especially leaf samples showed a very high rate of duplicated paired-end reads predominantly located in the chloroplast chromosomes [22]. To eliminate biases possibly arising from the high duplication rates, we performed a deduplication step using *samtools* markdup (version 1.9) [76] on each genomic mapping. Thus, all samples contained at least 30 million paired-end uniquely mapped deduplicated reads [22]. Subsequently, the mappings were curated in order to fix mating read pairs and to remove soft-clipped subsequences from the mapped reads.

The initial transcriptome assembly was performed using *cufflinks* (version 2.2.1) [55] with the parameters -b --library-type fr-firststrand --max-intron-length 2000 --min-isoform-fraction 0.1 --overlap-radius 1 --3- overhang-tolerance 1 --intron-overhang-tolerance 1. As input, we used the deduplicated mapped reads of each tissue or developmental stage represented by three biological replicates. A separate transcriptome assembly was performed for each biological replicate. Using cuffmerge [77], we summarized the assemblies of the three replicates into one organ-specific transcriptome assembly, resulting in eight organ-specific transcriptome assemblies for each plant species.

Next, the organ-specific assemblies were filtered by removing transcripts that spanned multiple loci or showed expression comparable to transcriptional noise. Using salmon [78], we quantified the expression abundance of each assembled transcript in each biological replicate. Transcriptional noise was defined as transcripts showing an expression level below the 5% quantile of expressed ERCC Spike-Ins in more than two out of three biological replicates. The organ-specific transcriptome assemblies and the reference annotation were combined using gffcompare (version 0.11.2) [79] into a species-specific transcriptome annotation.

The resulting transcriptome assembly was additionally filtered to remove transcripts showing insufficient transcript junction coverage. Therefore, we used GeMoMa (version 1.5.3) [80] with the parameters ERE c=true s=FR_FIRST_STRAND to calculate the splice junction coverage of each assembled transcript. We removed transcripts having a junction coverage of less than two split reads in at least two out of three biological replicates.

Subsequently, transcripts were clustered using CD-Hit (version 4.8.1) [81] with parameters −c 0.95 −n 8 −r 0 −p 1 −g 1 −d 0 to remove transcripts showing more than 95% sequence identity. The CD-Hit algorithm clustered the assembled transcripts, and the longest transcript in each cluster was selected as a representative while the remaining transcripts were discarded.

In addition, similarity searches using BLAST (version 2.9.0) [82] against the SILVA ribosomal RNA database [83] (retrieved 2019-02-15) were performed, and transcripts showing high similarity to ribosomal RNAs were removed. The resulting transcriptome assembly (Fig. 1a) served as the starting point for the annotation of novel protein-coding genes, alternative splice variants, long non-coding RNAs, and circular RNAs.

### Classification of coding and non-coding transcripts

The assembled transcripts were assigned into two groups (Supplementary Fig. 1). The first group contained all transcripts that overlap with known protein-coding loci. This group was further processed to annotate alternative splice variants. The second group contained all transcripts that showed no overlap with known protein-coding loci. For these transcripts, the coding potential was calculated to differentiate between candidate transcripts for putative novel protein-coding loci or lncRNAs. The coding potential was calculated using TransDecoder [30] (version 3.0.1) in combination with BLASTP (version 2.9.0) [82] searches against plant proteins from the Uniref90 database [84] and HMMER (version 3.1b2) [85] searches against the Pfam-A database [86]. The following main steps were performed:

1. Prediction of the longest open reading frame for each assembled transcript with unknown coding potential and translation into amino acid sequence by using TransDecoder.LongOrfs mod -S -t.
2. BLASTP searches of translated amino acid sequences from step 1 against plant proteins of the Uniref90 database using an e-value threshold of 1e-5.
3. HMMER searches with hmmscan of each amino acid sequence from step 1 against Pfam-A.
4. Training of TransDecoder [30] classifier with known species-specific coding sequences provided by the original genome annotation.
5. Classification of coding and non-coding transcripts based on the BLASTP and HMMER output, and the trained TransDecoder classifier using TransDecoder.Predict mod −t transcripts.fasta −−train knownCDS.fasta --retain_pfam_hits hmmer_res.domtblout --retain_blastp_hits blast_res.tblout --single_best_orf.

### Filtering putative novel protein-coding loci and alternative splice variants

We removed putative novel protein-coding transcripts overlapping with annotated transposable elements and *de novo* predicted transposable elements using LTRpred [87]. Next, we performed for each of the remaining putative coding transcripts an InterProScan [88] search against a variety of protein and domain sequence databases, e.g., CDD [89], Pfam [90], TIGRFAM [91], ProDom [92], to detect functional associations. The InterProScan 5 algorithm uses different search algorithms, e.g., BLAST [82] and HMMER [85]. We only considered transcripts that showed significant functional associations as novel protein-coding loci. For each potentially novel splice variant overlapping a known protein-coding gene, we calculated the pairwise global alignment [93] between the assembled transcript and each splice variant of the reference annotation at the overlapping gene locus. Subsequently, we predicted ORFs within the assembled transcript to ensure that the transcript was not truncated. We considered an assembled transcript as a new protein-coding splice variant only if it shows a minimum of 50% sequence identity with at least one reference splice variant and shared the same start codon position with its aligned target. Otherwise, the assembled transcript was regarded as incorrectly assembled and was discarded. The threshold of 50% sequence similarity is arbitrary and represents a compromise between predicting too many falsely assembled transcripts and removing too many putatively functional splice-variants. Subsequently, we filtered splice variants based on intron retention events. Since the RNA extraction protocol is intended to capture RNA species regardless of polyadenylation, it was likely to sequence unprocessed fragments of immature pre-mRNAs. It has been shown that total RNA sequencing results in a higher proportion of RNAs originating from intronic sequences [94, 95]. We assume that the assembled transcripts originated from intron retention might represent immature pre-mRNAs rather than functional mRNAs. To classify intron retention events, we used SUPPA (version 2.3) [96] with the parameters generateEvents −f ioe −e RI and removed all assembled transcripts showing retained introns.

### Prediction of long non-coding RNAs

Initially, we aimed to assign the annotated non-coding RNAs into sub groups of bidirectional, intergenic (lincRNA), exonic antisense (NAT), and intronic antisense (intronic NAT) lncRNAs (Supplementary Fig. 2a). The definition of lncRNA sub groups (Supplementary Fig. 2a) is based on observations in humans and other vertebrates [39]. Typically, lncRNAs in the proximity of protein-coding genes and also sharing the same promoter region are classified as bidirectional lncRNAs. This distance ranges from 500 nt to 5,000 nt [5, 20, 97], or up to 250 nt with respect to the smaller and more compact plant genomes in our study. However, compared to humans and other vertebrates a substantial absence of bidirectional transcription has been shown in *A. thaliana* seedlings [98]. Thus, despite the potential of classifying lncRNAs as bidirectional, we assigned lncRNAs in the proximity of protein-coding genes to the group of lincRNAs if they did not overlap with any exon of a protein-coding gene. Otherwise, we categorized them into the group of NATs (Supplementary Fig. 2a).

As described earlier, total RNA-Seq data attends to be biased by an overrepresentation of reads in intronic regions [29]. Thus, we discarded all candidate intronic sense lncRNA transcripts because we were not able to distinguish intronic (sense) transcripts from sequencing or RNA extraction artifacts. Additionally, we eliminated non-coding transcripts overlapping with any annotated or predicted transposable element.

By using TransDecoder [30] to classify coding and non-coding transcripts, we were able to train the underlying algorithm on known protein-coding transcripts for each reference. To validate our procedure, we compared the TransDecoder results with popular lncRNA prediction tools such as FEELnc [99], which can be trained only on known species-specific coding transcripts, and CPC2 [100], which is already trained on sets of non-coding and protein-coding transcripts. We compared the accuracy of non-coding predictions between FEELnc [99], CPC2 [100], and TransDecoder [30] using a training dataset [100]. CPC2 and TransDecoder achieved a prediction accuracy of ∼97% for *A. thaliana*, while FEELnc showed an accuracy of ⇠94%, confirming our choice of using TransDecoder (Supplementary Fig. 2b).

### Prediction of circular RNAs

The detection of circular RNAs (circRNAs) was performed on the updated transcriptome annotation, which already included the newly assembled alternative splice variants, putative protein-coding loci, and lncRNAs. We used DCC [101] with the parameters -D -Pi -F -Nr 1 2 -fg -G to annotate novel circRNAs. According to the parameter “-Nr 1 2”, a circRNA was identified only if it could be predicted in at least two biological replicates, each covered at least by one back-spliced read (Supplementary Fig. 3a).

To evaluate the performance of DCC, we benchmarked the most common circRNA prediction tools such as CIRI2 [102], circExplorer2 [103], and DCC [101]. These tools have already been evaluated in various studies, but mainly focusing on human or other animal data sets [101, 104].

The benchmark was based on our 132 the sequencing libraries of *A. thaliana* covering 10 organs and different developmental stages [22]. We compared our predictions with already published and annotated circRNAs provided by the PlantcircBase database [105]. For *A. thaliana*, the database contained 38,938 circRNAs (version 4, 2019), of which 29,348 had strand information and were included in our analysis (Supplementary Fig. 3b).

---

**Supplementary Figure 1.**
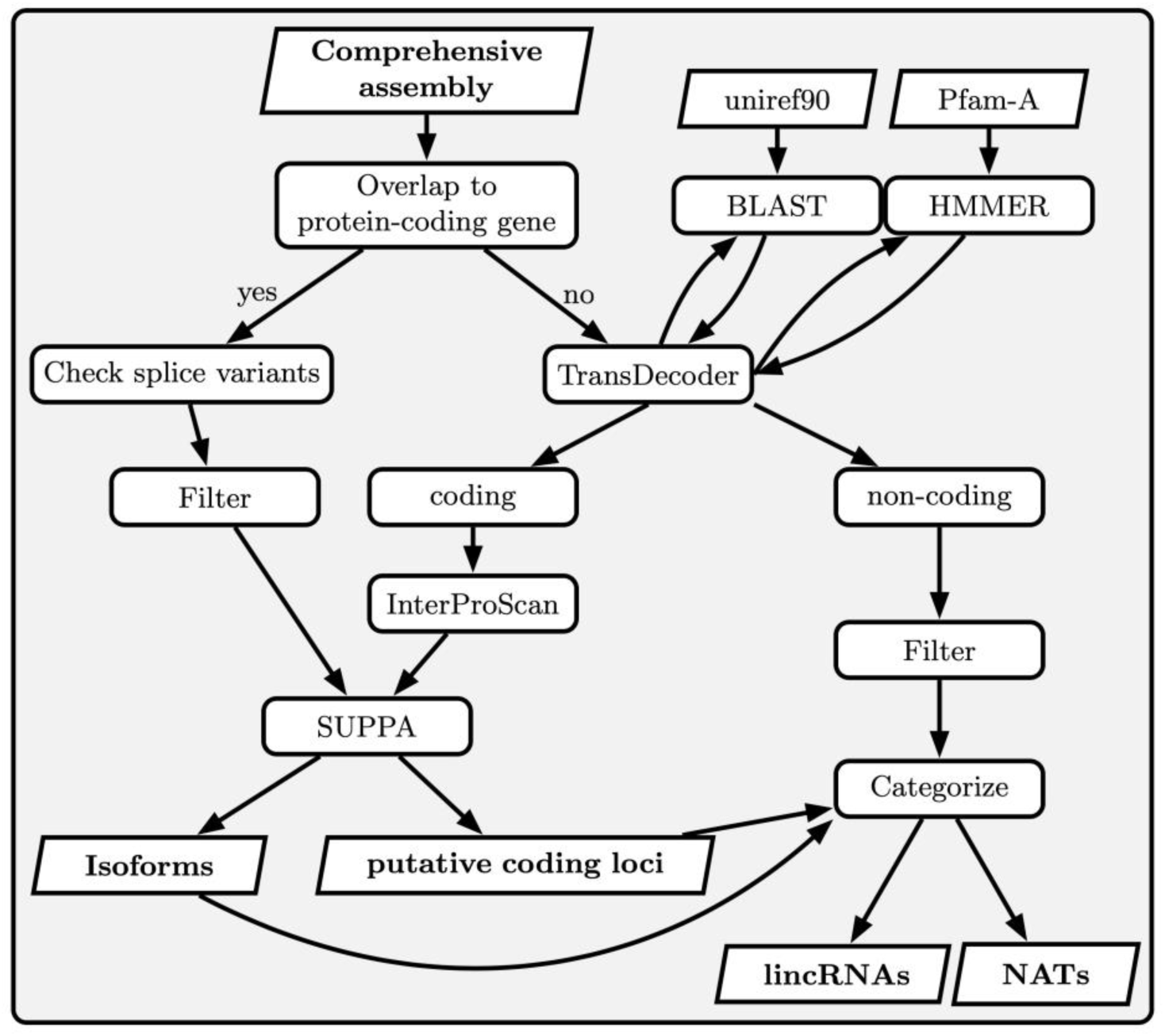
annotation of protein-coding and non-coding transcripts. Starting from the comprehensive assembly, each transcript is first assigned to known protein-coding transcripts or unknown transcripts. The known transcripts were filtered and used to update the known protein-coding transcriptome with novel isoforms or replace known isoforms. Unknown transcripts were assigned into potentially coding, i.e., probably putative protein-coding loci, and non-coding. The non-coding transcripts were filtered and categorized as intergenic or antisense lncRNA (NAT).

**Supplementary Figure 2.**
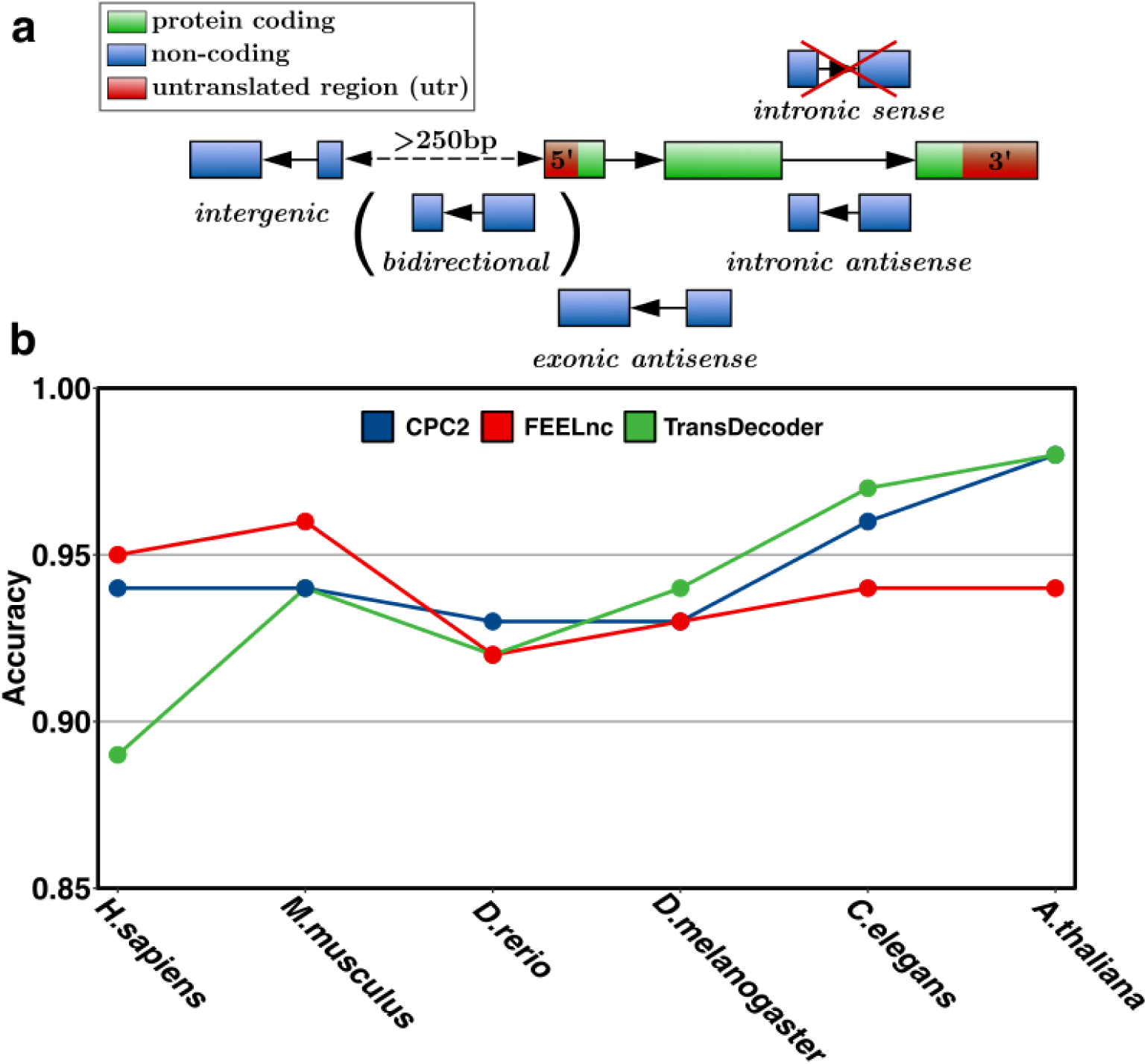
Classification of lncRNAs. a. The quantification of lncRNAs (blue) is based on their transcript length (>200nt), and its genomic location relative to a proximal protein-coding gene (green and red). In theory lncRNA transcripts can be divided into five distinct classes: (i) intergenic lncRNAs (lincRNAs), (ii) bidirectional lncRNAs, (iii) exonic antisense lncRNAs, (iv) and intronic antisense lncRNAs (NATs), and (v) intronic sense lncRNAs. Our RNAseq data was not sufficient to reliably predict intronic sense lncRNAs due to the fact that library preparation of total RNAs leads to a systematic bias of generating immature spliced transcripts, which often leads to an increased probability of falsely discovering intronic sense lncRNAs. Bidirectional lncRNAs (here shown in brackets) were counted as intergenic lncRNAs due to the lack of bidirectional promoters in plant species. b. Comparative assessment of computational tools specialized in detecting and quantifying the coding potential of transcripts. Here, we assess the accuracy of each tool in the context of reliably predicting lncRNAs. Notably, plant lncRNAs are most sufficiently captured by TransDecoder and CPC2 which illustrated the highest accuracy (∼97%).

**Supplementary Figure 3.**
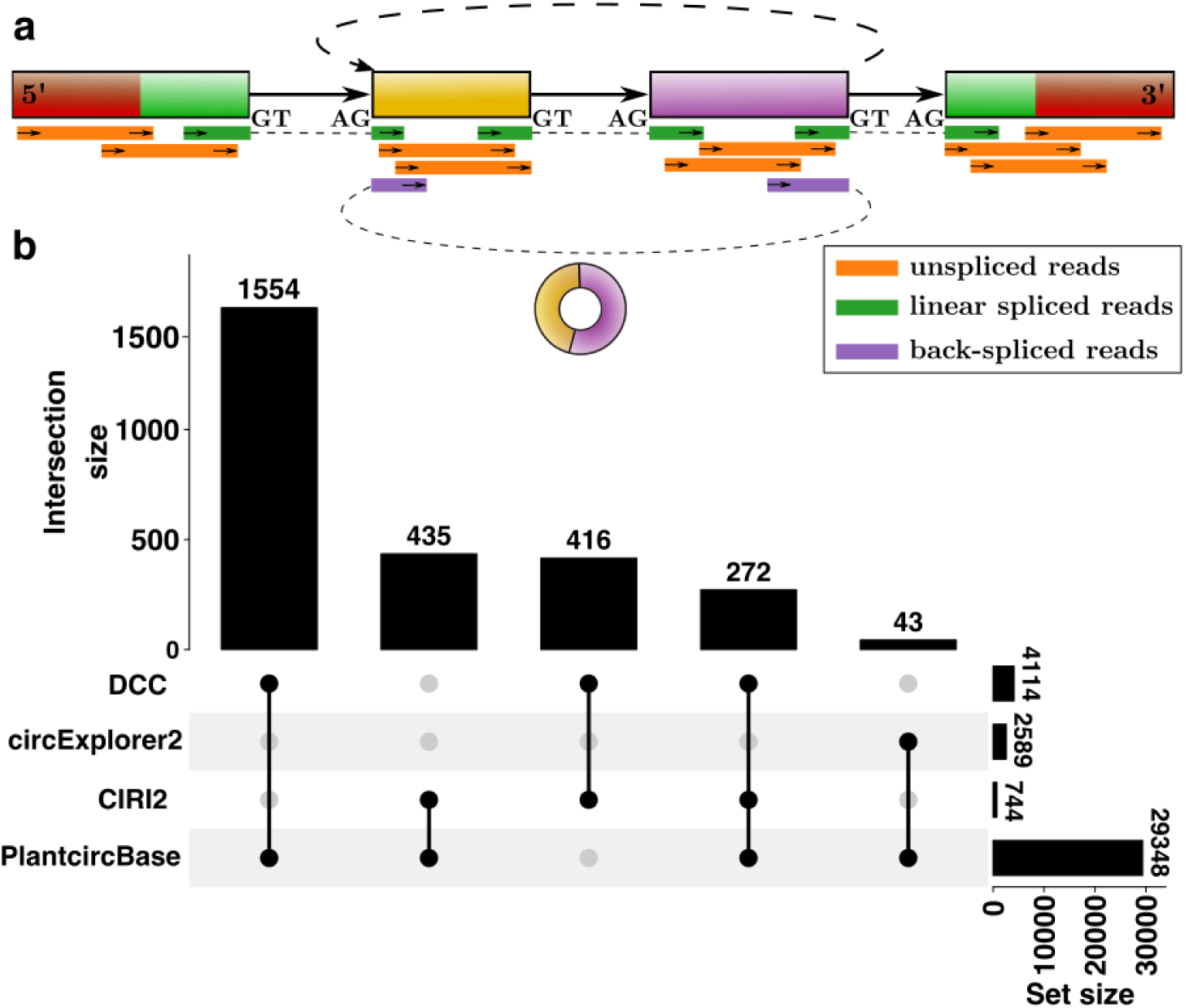
annotation of circRNAs in *A. thaliana*. a. Prediction of circRNAs within protein-coding genes of *A. thaliana*. Using back-spliced reads (violet), prediction tools are able to annotate circRNAs. In our analysis at least two biological replicates should provide evidence of a circRNA that based on back-spliced reads. b. Comparison of circRNA prediction tools. In total, DCC predicts the most circRNAs but also the most circRNAs which overlap with published circRNAs in the PlantcircBase [105].

**Supplementary Figure 4.**
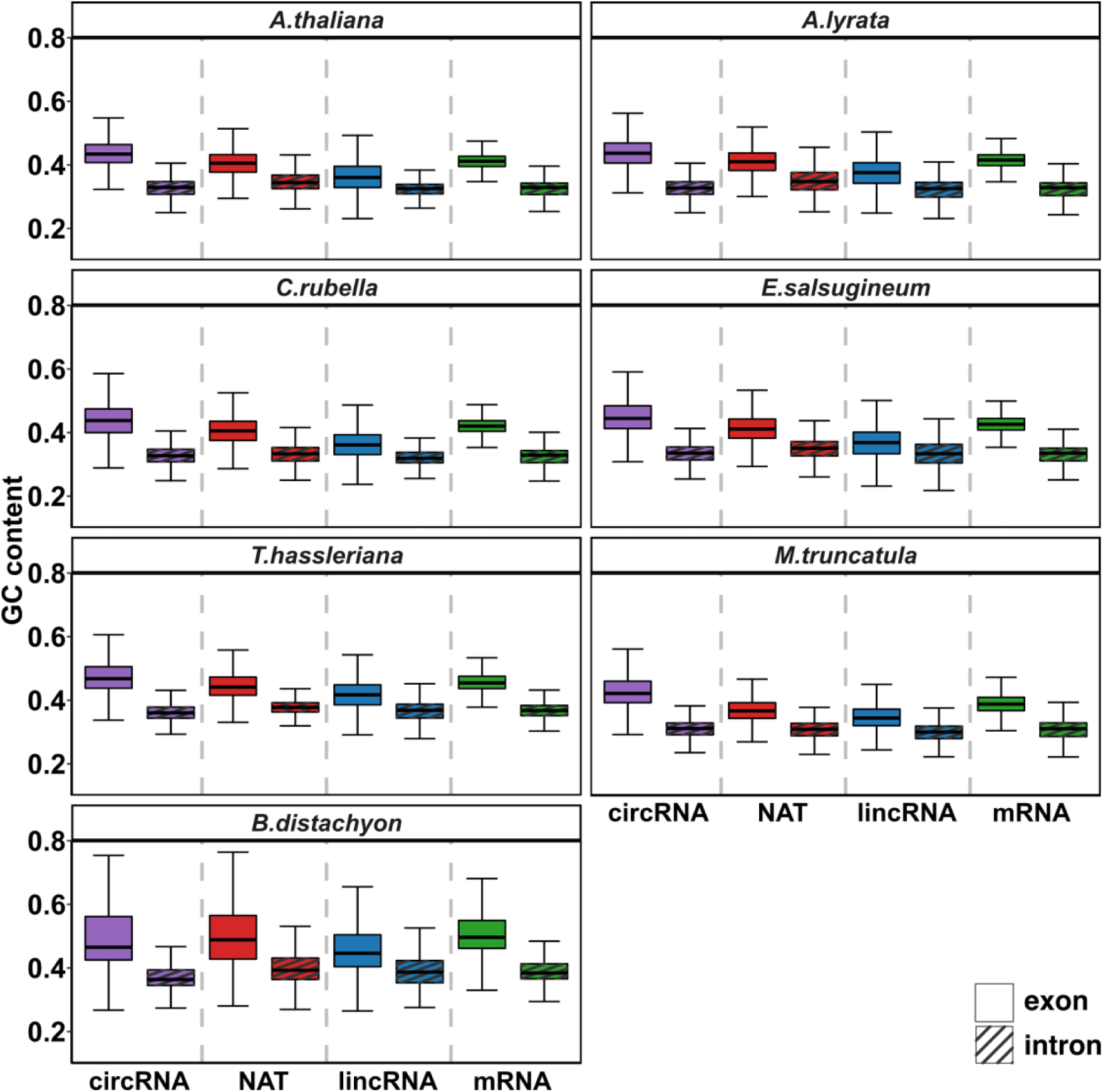
GC content in exons *versus* introns compared across different RNA species. For each plant species, we calculated the relative GC content for both exons and introns of circRNAs, lincRNAs, NATs, and protein-coding mRNAs. Exonic sequences tend to exhibit higher GC content and a broader IQR compared to intron derived sequences. For the majority of plant species, the range of GC content of the different RNA species is similar, except *B. distachyon*. *B. distachyon* shows the largest proportion of GC across the seven plant species and the widest IQR across all RNA species.

## Supplementary Tables (Supplementary_tables_devseq_annotation.xlsx)

**Supplementary Table 1**. BUSCO evaluation of annotation completeness. Absolute number (and percentage) of identified embryophyta specific single copy orthologs in BUSCO.

**Supplementary Table 2**. Novel putative protein-coding loci. Comparison of known protein-coding loci in the reference annotation, novel putative protein-coding loci, and the corresponding transcript isoforms identified by the DevSeq workflow.

**Supplementary Table 3**. Statistics of protein-coding isoforms. The columns “known” and “new” show the numbers of protein-coding isoforms in the reference annotation and the novel transcripts from the DevSeq workflow. Besides the prediction of novel isoforms, we updated known isoforms by elongating their 3’ and/or 5’ ends.

**Supplementary Table 4.** Sources of reference annotations. List of reference annotations and their corresponding genome sequences used by the DevSeq workflow.

**Supplementary Table 5**. GC content and quintiles of transcript lengths in coding and non-coding RNA species. For each species and each RNA species (except intronic NATs), we group the corresponding transcripts into quintiles based on their transcript length in nt and calculate the relative mean GC content ± standard deviation. The transcript length for each quanitle is given as an interval representing the minimal and maximal transcript length.

